# First-generation EGFR-TKI plus chemotherapy versus EGFR-TKI alone as first-line treatment in advanced NSCLC with EGFR activating mutation: a systematic review and meta-analysis of randomized controlled trials

**DOI:** 10.1101/2020.04.17.046409

**Authors:** Qiang Wu, Wuxia Luo, Wen Li, Ting Wang, Lin Huang, Feng Xu

## Abstract

The aim of this meta-analysis was to evaluate efficacy and toxicity of epidermal growth factor receptor tyrosine kinase inhibitor (EGFR-TKI) in combination with chemotherapy (CT) compared to EGFR-TKI monotherapy as first-line treatment in advanced non-small cell lung cancer (NSCLC) harboring activating EGFR mutation. A systematic literature search of randomized controlled trials using Cochrane Library, PubMed, Embase, and Web of Science, was performed up to Jan. 7^th^, 2020. A total of eight randomized trials involving 1,349 advanced NSCLC patients with sensitive EGFR mutation were included in the meta-analysis. All patients in both groups received first-generation TKI as first-line treatment. The pooled HR of PFS and OS was 0.56 (95% CI=0.50-0.64; P < 0.00001) and 0.70 (95% CI=0.54-0.90; P=0.005), respectively. Subgroup analysis showed significantly higher OS advantages in patients receiving doublet CT (P=0.02) and concurrent therapy (P=0.002). The ORR in the EGFR-TKI plus CT group was significantly higher than in the EGFR-TKI monotherapy group (RR = 1.18, 95% CI = 1.10–1.26). The combination regimen showed a higher incidence of chemotherapy-induced toxicities. Subgroup analysis indicated that doublet chemotherapy rather than single-agent chemotherapy significantly increased incidence of grade 3 or higher leukopenia, neutropenia and anemia. In conclusion, compared with EGFR-TKI monotherapy, the combination of first-generation EGFR-TKI and CT, especially when applying concurrent delivery of platinum-based doublet chemotherapeutic drugs, significantly improve ORR and prolong PFS and OS in first-line treatment for advanced EGFR-mutated NSCLC, yet at the price of increasing incidence of chemotherapy-induced toxicities that is well tolerated and clinically manageable.

## Introduction

Lung cancer is the leading cause of cancer morbidity and mortality worldwide, with 2.1 million new cases and 1.8 million deaths estimated in 2018 [1]. Non-small cell lung cancer (NSCLC) accounts for nearly 85% of all cases of lung cancer. Due to ineffective screening method and insidious symptom, lung cancer is usually diagnosed at an advanced stage in a majority of patients. Systematic therapy, therefore, remains the pivotal treatment approach for NSCLC in clinical practice.

Epidermal growth factor receptor (EGFR) is one of the most significant driver gene in lung cancer and its mutated form tempts constitutive activation of the EGFR tyrosine kinase, leading to uncontrolled growth and proliferation of tumor. Approximately, 10–15% of NSCLC patients in Europe and 30–35% of NSCLC patients in Asia harbor activating EGFR mutation [2, 3]. An Individual Patient Data Meta-Analysis of six large randomized controlled trials (RCTs) suggested that compared with chemotherapy, first-line EGFR tyrosine kinase inhibitor (TKI) significantly prolonged progression-free survival (PFS) (median PFS = 11.0 vs. 5.6 months; Hazard Ratio (HR) = 0.37, 95% confidence intervals (CI) = 0.32-0.42, P < 0.001) in EGFR-mutated NSCLC patients [4]. Thus, first-line EGFR-TKI monotherapy, including representative gefitinib and erlotinib, is currently the mainstay treatment for naive advanced EGFR mutation positive NSCLC patients [5].

Inevitably, most patients who initially respond to an EGFR-TKI over 8-12 moths, eventually develop resistance to first- or second-generation drugs [6]. In order to prolong the survival outcome, combination therapy of EGFR-TKI with other therapeutic drugs is an emerging promising approach. As one of promising combined strategy, EGFR-TKI plus chemotherapy has long been evaluated to overcome or delay resistance in advanced NSCLC since the early 2000s [7]. Due to lack of EGFR-mutation status selection, however, preliminary studies failed to demonstrate the survival benefit of EGFR-TKI in combination with chemotherapy [8, 9]. Recently, many phase II-III RCTs have investigated the EGFR-TKI plus chemotherapy combination in selected NSCLC patient with activating EGFR mutation [10]. These studies with EGFR sensitive mutation had mixed overall survival (OS) results. Meta-analysis assessing the efficacy and toxicity of EGFR-TKI in combination with chemotherapy as first-line treatment in advanced NSCLC with EGFR activating mutation, has not yet been reported to our best knowledge. Therefore, we synthesized the results of different studies in this meta-analysis, to provide more objective data for the optimal clinical use of EGFR-TKI combined with chemotherapy.

## Material and Methods

### Search strategy

Our study was performed in accordance with the Preferred Reporting Items for Systematic Reviews and Meta-Analyses (PRISMA) statement [11]. A comprehensive search of PubMed, Embase, Web of Science, and Cochrane databases was conducted to identify all relevant full-length literatures on the comparison of EGFR-TKI plus chemotherapy to EGFR-TKI alone as first-line treatment in advanced non-small cell lung cancer with activating EGFR mutation, up till Jan. 7^th^, 2020. Keywords including non-small-cell lung cancer, EGFR, TKI, and chemotherapy were used for initial search of eligible literatures. For instance, the following retrieval strategy was used on PubMed: (lung cancer OR lung carcinoma OR lung neoplasm) AND (epidermal growth factor receptor OR EGFR) AND (tyrosine kinase inhibitor OR TKI OR gefitinib OR erlotinib OR icotinib OR afatinib OR dacomitinib OR osimertinib) AND (chemotherapy OR pemetrexed OR gemcitabine OR paclitaxel OR vinorelbine) AND (first line OR untreat* OR naive). To obtain additional related articles, references cited in the eligible studies were also searched manually.

### Selection criteria

Inclusion criteria were as follows: (1) the patients were histologically diagnosed with advanced NSCLC; (2) the randomized trials were performed to evaluate the efficacy and safety of compared the EGFR-TKI plus chemotherapy to EGFR-TKI alone as first-line treatment in advanced non-small cell lung cancer with activating EGFR mutation; (3) the studies with affluent data for pooling the survival results, response rate, and toxicity. Exclusion criteria were as follows: (a) nonoriginal research articles with limited data, such as letters, case reports, reviews, comments, and conference abstracts; (b) duplicates of previous publications; and (c) studies with a sample size of less than 30 analyzable lesions.

### Data extraction and quality assessment

Basic information of each individual study was extracted by two reviewers (QW and WXL) independently. Any discrepancies were resolved by discussion and consensus during the process of research selection and data extraction or by consulting the third investigator (FX) when necessary. The following information was extracted: name of first author, trial name, publication year, trial phase, treatment arms, participants’ characteristics, number of patients evaluable for analysis and other clinical characteristics. The primary data for calculation were the HR with 95% CI for PFS and OS, the number of patients who experienced a partial response or complete response, the number of patients that developed all grade toxicities.

A specific tool recommended by the Cochrane Collaboration was applied to assess the risk of bias for each identified study. Biases were categorized as selection bias, performance bias, detection bias, attrition bias, and reporting bias [12].

### Statistical analysis

All statistical analysis was performed using the Review Manager 5.3 software (Cochrane Library, Oxford, UK) and STATA 12.0 software (Stata Corp., College Station, TX). Cochrane’s Q statistic and I^²^ (I^2^>50% was considered substantially heterogeneous) statistic test were used to evaluate the heterogeneity between the eligible studies [13]. The random effect model was used when there was significant heterogeneity between studies; otherwise, the fixed effect model was used. Publication bias was assessed via funnel plot with Begg’s rank correlation. A two-sided p-value of <0.05 was considered to be statistically significant.

## Results

### Study selection

A comprehensive literature search yielded 1732 non-duplicate papers. Of these, 21 full-text articles were screened for assessment of eligibility in the review. Of these potential studies, 13 studies were excluded because of retrospective studies [14–17], unselected EGFR mutation [18–21], single arms [22–24], small sample size [25], and insufficient survival data [26]. Finally, eight studies comparing EGFR-TKI plus chemotherapy with EGFR-TKI alone as first-line treatment in advanced non-small cell lung cancer with activating EGFR mutation were included in this meta-analysis [8, 27–33]. The flow diagram of studies selection was summarized in Fig 1.

**Fig 1.**
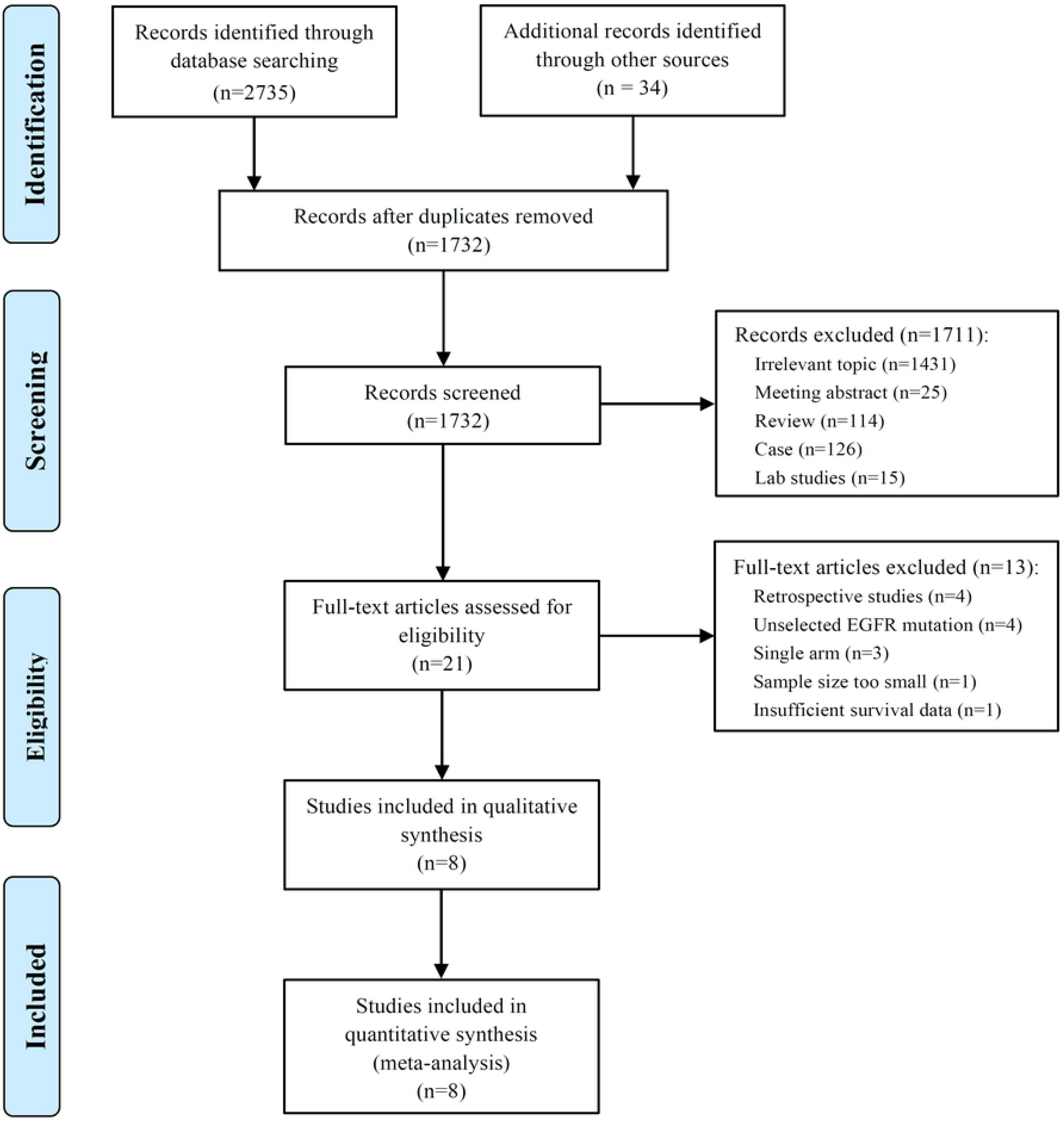
The flow diagram of studies selection.

### Characteristics of eligible studies

From eight clinical trials, a total of 1,349 advanced NSCLC patients with sensitive EGFR mutation (704 in EGFR-TKI combination group and 645 in EGFR-TKI monotherapy group), were available for the meta-analysis. The great majority of histological type was adenocarcinoma. Of these EGFR-mutated patients, exon 19 deletion and L858R point mutation accounted for 55.7% (751/1,349) and 40.9% (552/1,349), respectively. As for first-line EGFR-TKI treatment, patients in all trials received first-generation drugs, including gefitinib (six studies), erlotinib (one study) and icotinib (one study). Most of trials involved platinum-based doublet chemotherapy, apart from two trials [8, 29]. In addition, concurrent drug delivery of TKI and chemotherapy were engaged in four of these studies. The characteristics of the included studies are listed in Table 1.

**Table 1.** Charactetistics of the included randomized trials in the meta-analysis

| Study | Year | Country | Phase | Group | Type of combination | No. of evaluable patients | Median age (years) | No. of EGFR mutation |  | Adenocarcinoma (%) | Efficacy |  |  |
| --- | --- | --- | --- | --- | --- | --- | --- | --- | --- | --- | --- | --- | --- |
|  |  |  |  |  |  |  |  | 19 del | L858R |  | ORR | PFS | OS |
| CALGB 30406 | 2012 | USA | II | Paclitaxel plus carboplatin+ E | Concurrent | 33 | 60 | 16 | 17 | 84 | 73% | 17.2 m | 38.1 m |
|  |  |  |  | E |  | 33 | 58 | 23 | 10 | 88 | 70% | 14.1 m | 31.3 m |
| Yang | 2014 | East Asia | III | Pemetrexed plus cisplatin +G | Sequential | 26 | 59 | 14 | 10 | 97 | 65.4% | 12.9 m | 32.4 m |
|  |  |  |  | G |  | 24 | 59 | 11 | 13 | 97 | 70.8% | 16.6 m | 45.7 m |
| An | 2016 | China | II | Pemetrexed +G | Intercalated | 45 | 65.7 | 16 | 29 | 100 | 80.0% | 18.0 m | 34.0 m |
|  |  |  |  | G |  | 45 | 66.9 | 17 | 28 | 100 | 73.3% | 14.0 m | 32.0 m |
| Cheng | 2016 | East Asia | II | Pemetrexed +G | Concurrent | 126 | 62 | 65 | 52 | NA | 80.2% | 15.8 m | 43.4 m |
|  |  |  |  | G |  | 65 | 62 | 40 | 23 | NA | 73.8% | 10.9 m | 36.8 m |
| Han | 2017 | China | II | Pemetrexed plus carboplatin +G | Intercalated | 40 | NA | 21 | 19 | 100 | 82.5% | 17.5 m | 32.6 m |
|  |  |  |  | G |  | 41 | NA | 21 | 20 | 100 | 65.9% | 11.9 m | 25.8 m |
| NEJ009 | 2019 | Japan | III | Pemetrexed plus carboplatin +G | Concurrent | 170 | 64.8 | 93 | 69 | 98.8 | 84% | 20.9 m | 50.9 m |
|  |  |  |  | G |  | 172 | 64.0 | 95 | 67 | 98.8 | 67% | 11.9 m | 38.8 m |
| Noronha | 2019 | India | III | Pemetrexed plus carboplatin +G | Concurrent | 174 | 54 | 107 | 60 | 98 | 75.3% | 16.0 m | NR |
|  |  |  |  | G |  | 176 | 56 | 109 | 60 | 97 | 62.5% | 8.0 m | 17.0 m |
| Xu | 2019 | China | II | Pemetrexed plus carboplatin +I | Intercalated | 90 | 58.6 | 51 | 38 | 100 | 77.8% | 16.0 m | 36.0 m |
|  |  |  |  | I |  | 89 | 61.0 | 52 | 37 | 100 | 64.0% | 10.0 m | 34.0 m |

E: erlotinib; G: Gefitinib; I: icotinib; ORR: objective response rate; PFS: progression free survival; OS: over survival; NA: not available; NR: not reach; EGFR: epidermal growth factor receptor; m: months

### Progression-free survival

The median PFS as the primary end point of the studies ranged from 7.2 to 20.9 months in the EGFR-TKI combination arm and ranged from 4.7 to 16.6 months in EGFR-TKI monotherapy arm. The heterogeneity was not significant (I^2^=11%; P =0.34), and hence a fixed-effects model was used to pool the data (Fig 2A). The pooled HR of PFS in total population with activating EGFR mutation was 0.56 (95% CI=0.50-0.64; P < 0.00001; Fig 2A), which indicated that EGFR-TKI combination therapy significantly reduced the risk of disease progression compared with EGFR-TKIs alone. Furthermore, the pooled HR of PFS in patients with Exon 19 deletion or L858R point mutation was 0.54 (95% CI=0.45-0.65; P < 0.00001; Fig 2B) and 0.52 (95% CI=0.42-0.65; P < 0.00001; Fig 2C), respectively, retrieved from five included studies.

**Fig 2.**
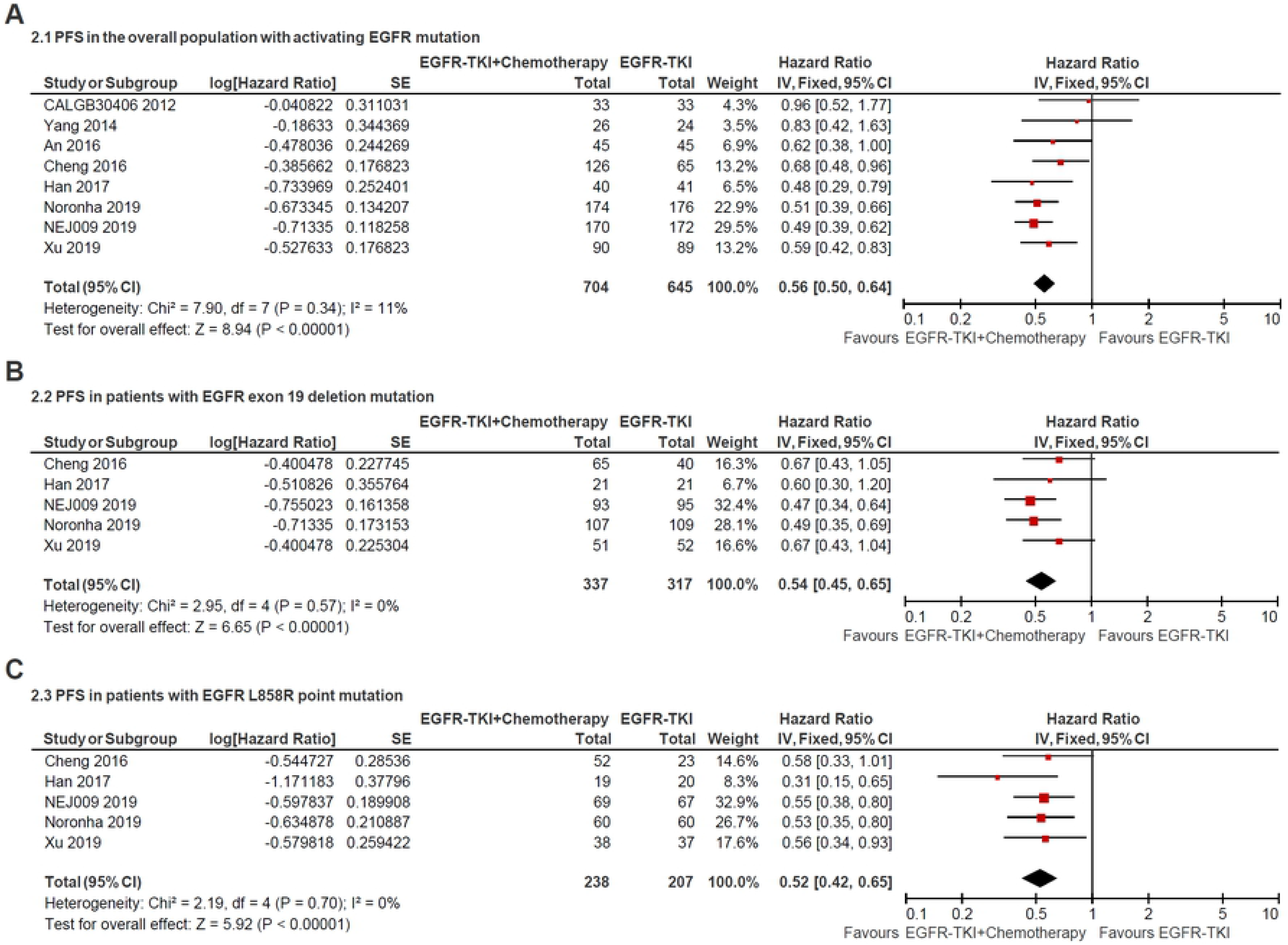
Forest plot of hazard ratio of progress-free survival in overall patients with all sites of positive activating EGFR mutation (A); in patients with positive exon 19 deletion mutation (B) and positive L858R point mutation (C).

Subgroup analysis of chemotherapy drugs revealed that double-agents mighty induced more longer PFS (double-agents, HR=0.54, 95% CI=0.47-0.62; single-agent, HR=0.66, 95% CI=0.50-0.87; Fig 3A). Moreover, subgroup analysis of combination timing indicated statistically significant PFS in concurrent and intercalated therapy (HR=0.55, 95% CI=0.47-0.64 and HR=0.57, 95% CI=0.45-0.73, respectively; Fig 3A), but was not statistically significant in sequential therapy (HR=0.83; 95% CI= 0.42–1.63; Fig 3A).

**Fig 3.**
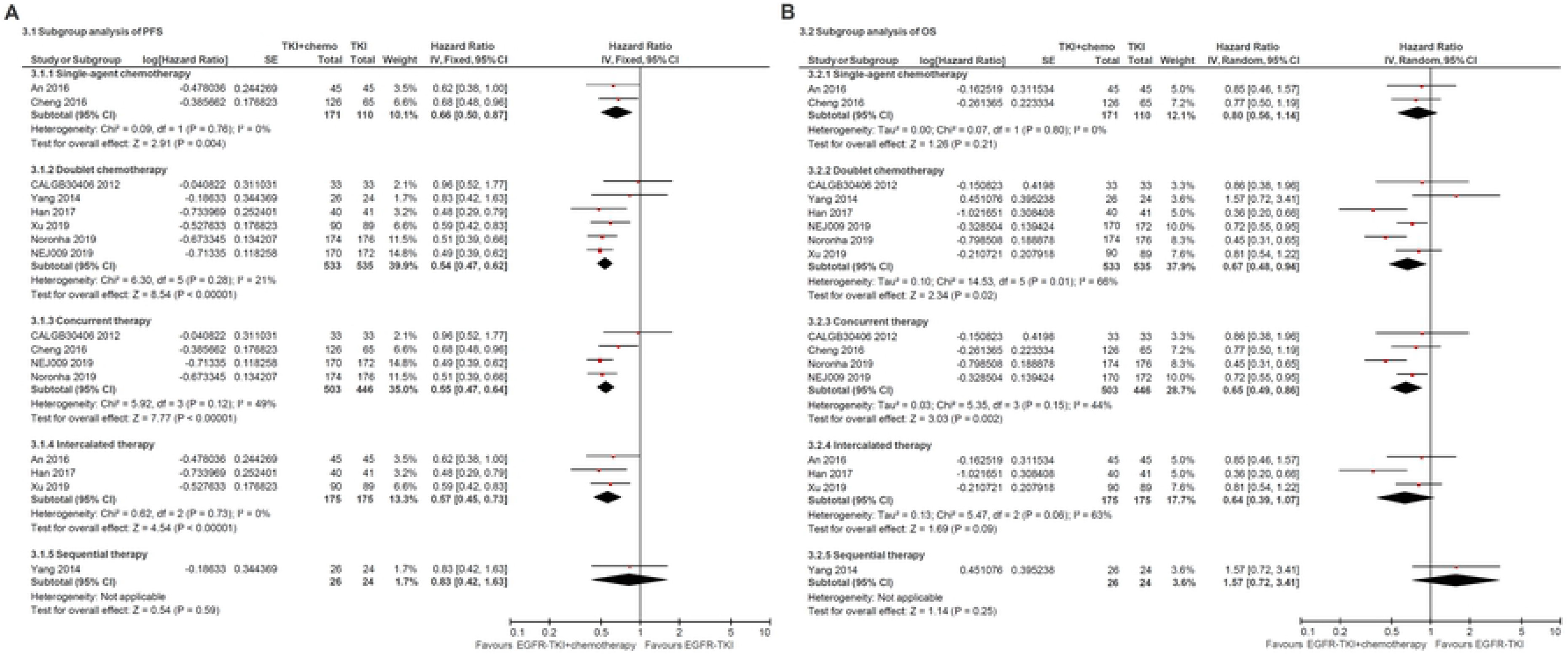
Forest plot of subgroup analysis of progress-free survival (A) and overall survival (B).

### Overall survival

The median OS in the included studies ranged from 18.5 to 50.9 months in the EGFR-TKI combination arm and ranged from 14.2 to 45.7 months in EGFR-TKI monotherapy arm. The pooled HR of OS in total EGFR sensitive mutation sites between two arms was 0.70 (95% CI=0.54-0.90; P=0.005; Fig 4A), which indicated that combination therapy significantly improved the OS compared with EGFR-TKIs alone. Furthermore, the pooled HR of OS in patients with exon 19 deletion or L858R point mutation was 0.60 (95% CI=0.42-0.86; P=0.005; Fig 4B) and 0.82 (95% CI= 0.57-1.18; P=0.28), respectively, retrieved from two trials. It revealed that overall survival benefit from EGFR-TKI in combination with chemotherapy might occur in patients with positive 19 deletion mutation other than in L858R mutation.

**Fig 4.**
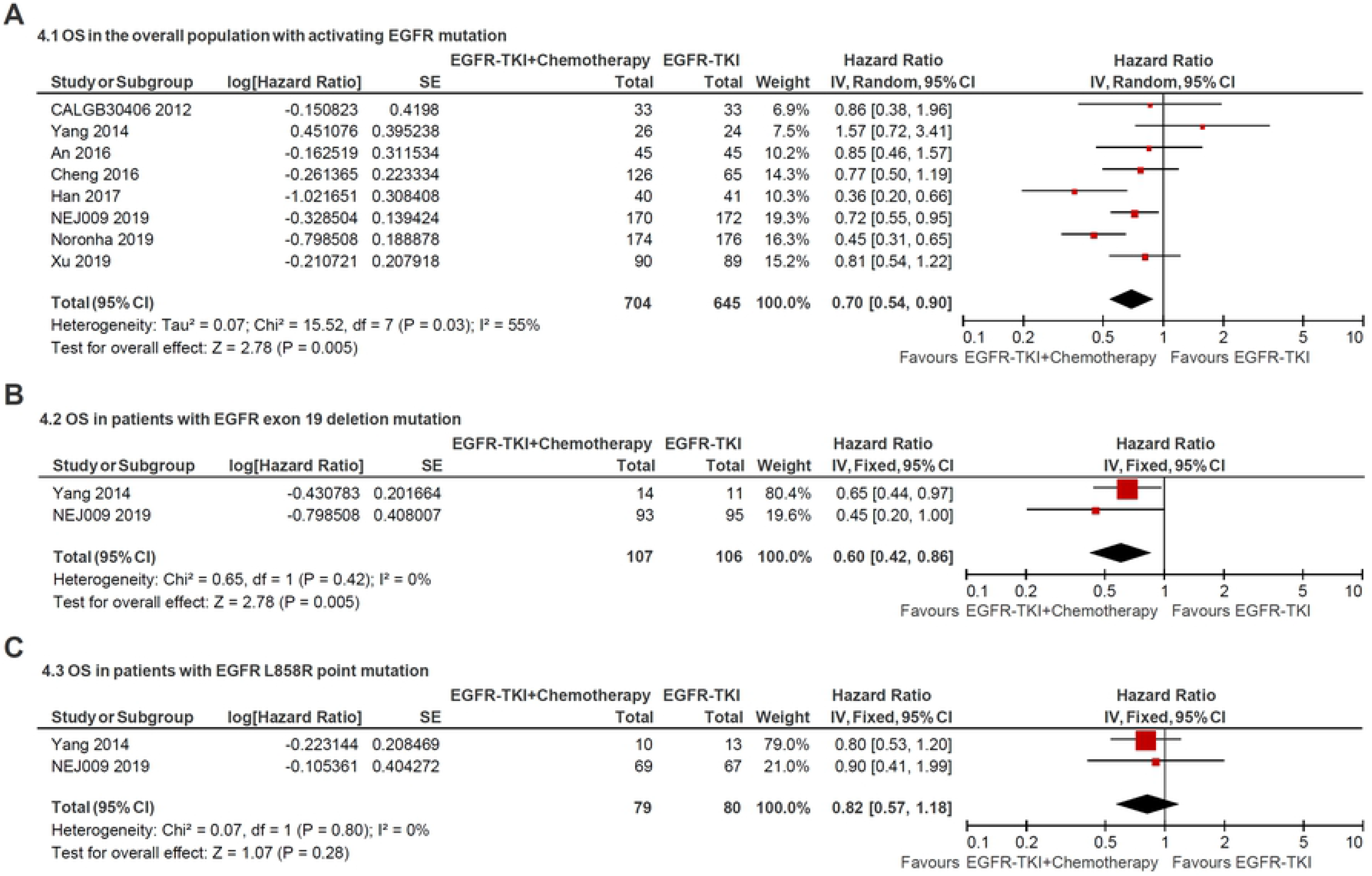
Forest plot of hazard ratio of overall survival in patients with all sites of positive activating EGFR mutation (A); in patients with positive exon 19 deletion mutation (B) and positive L858R point mutation (C).

Subgroup analysis of chemotherapy drugs revealed that double-agents significantly improved PFS (double-agents, HR=0.67, 95% CI=0.48-0.94; single-agent, HR=0.80, 95% CI=0.56-1.14; Fig 3B). In addition, subgroup analysis of combination timing indicated statistically significant OS in concurrent therapy (HR=0.65, 95% CI=0.49-0.86; Fig 3B), but was not statistically significant in intercalated and sequential therapy (HR=0.64, 95% CI=0.39-1.07 and HR=1.57, 95% CI=0.72-3.41, respectively; Fig 3B).

### Objective response rate

All of eight studies reported the data of objective response rate (ORR). The heterogeneity was not significant (I^2^ = 0%; P = 0.66), and hence a fixed-effects model was used to pool the data (Fig 5). The meta-analysis demonstrated that pooled ORR in the EGFR-TKI plus chemotherapy group was significantly higher than in the EGFR-TKI monotherapy group (RR = 1.18, 95% CI = 1.10–1.26; p < 0.00001; Fig 5).

**Fig 5.**
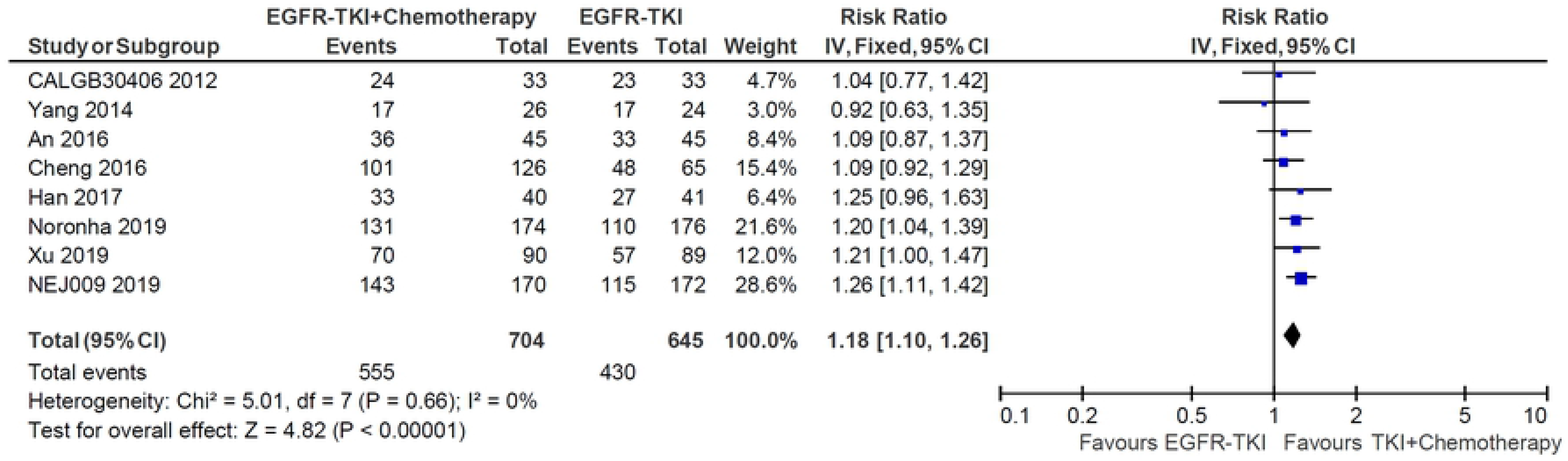
Forest plot of Risk ratio of objective response rate in EGFR-TKI plus chemotherapy group and EGFR-TKI monotherapy group.

### Toxicities

Compared with the EGFR-TKI alone, the addition of chemotherapy to TKI was associated with a higher incidence of any grade hematologic toxicities, such as neutropenia (grade 1-2, OR=16.84, 95% CI=9.04-31.36; grade 3 or higher, OR=10.03, 95% CI=1.04-96.69) and thrombocytopenia (grade 1-2, OR=7.04, 95% CI=4.73-10.48; grade 3 or higher, OR=43.41, 95% CI=6.01-313.71). Similarly, the combination therapy significantly increased the incidence of chemotherapy-induced toxicities, including any grade fatigue, anorexia, nausea and vomiting, and grade 3 or higher diarrhea. Subgroup analysis indicated that doublet chemotherapy significantly increased incidence of grade 3 or higher leukopenia (OR=37.30, 95% CI=7.26-191.63, Fig 6A), neutropenia (OR=54.79, 95% CI=13.21-227.24, Fig 6B) and anemia (OR=14.28, 95% CI=6.10-33.43, Fig 6C), while those differences were not significant in single-agent chemotherapy subgroup.

**Fig 6.**
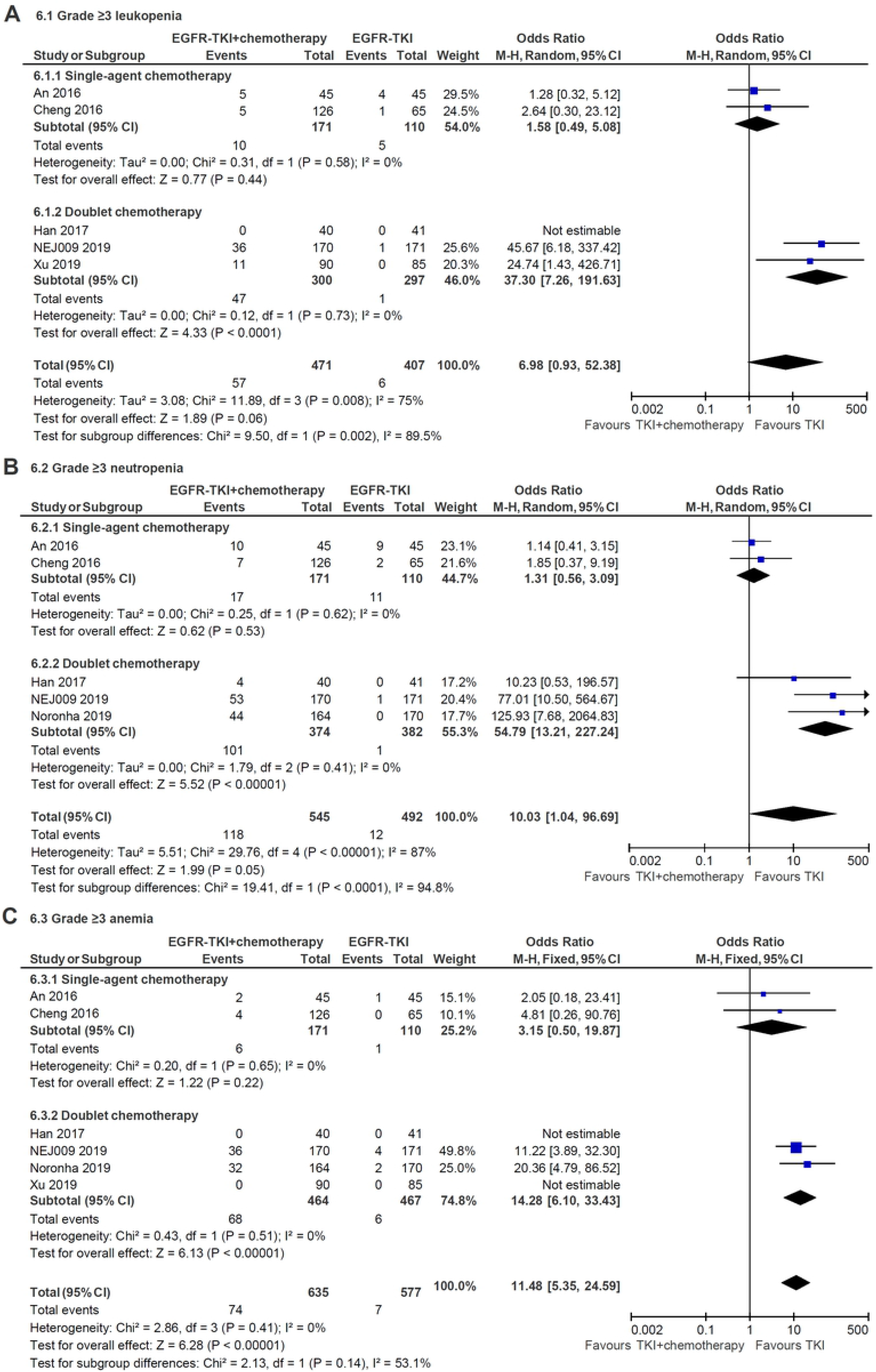
Subgroup analysis of grade 3 or higher hematologic toxicities for leukopenia (A), neutropenia (B) and anemia (C) in single-agent and doublet chemotherapy.

Nevertheless, no significant differences were founded in terms of any grade rash and grade 3 or higher liver dysfunction when applying combined treatment. The detail results are illustrated in Table 2.

**Table 2.** Pooled odds ratio of toxicities in included randomized trials

| Toxicities | No. of trials | Events in EGFR-TKI+Chemotherapy group | Events in EGFR-TKI monotherapy group | Odds Ratio (95% CI) | P value | Heterogeneity (I<sup>2</sup>) (%) |
| --- | --- | --- | --- | --- | --- | --- |
| Leukopenia |  |  |  |  |  |  |
| Grade 1-2 | 4 | 117/426 | 24/362 | 5.72 (3.59-9.12) | < 0.001 | 45 |
| Grade ≥ 3 | 5 | 57/471 | 6/407 | 6.98 (0.93-52.38) | 0.06 | 75 |
| Neutropenia |  |  |  |  |  |  |
| Grade 1-2 | 4 | 143/500 | 11/447 | 16.84 (9.04-31.36) | < 0.001 | 48 |
| Grade ≥ 3 | 5 | 118/545 | 12/492 | 10.03 (1.04-96.69) | 0.05 | 87 |
| Thrombocytopenia |  |  |  |  |  |  |
| Grade 1-2 | 4 | 164/464 | 39/467 | 7.04 (4.73-10.48) | < 0.001 | 0 |
| Grade ≥ 3 | 5 | 37/509 | 0/512 | 43.41 (6.01-313.71) | < 0.001 | 0 |
| Anemia |  |  |  |  |  |  |
| Grade 1-2 | 5 | 261/590 | 158/532 | 3.01 (1.72-5.28) | < 0.001 | 58 |
| Grade ≥ 3 | 6 | 74/635 | 7/577 | 11.48 (5.35-24.59) | < 0.001 | 0 |
| Liver dysfunction |  |  |  |  |  |  |
| Grade 1-2 | 5 | 308/590 | 206/532 | 1.71 (1.34-2.19) | < 0.001 | 0 |
| Grade ≥ 3 | 6 | 82/635 | 63/577 | 1.56 (0.74-3.31) | 0.25 | 68 |
| Rash |  |  |  |  |  |  |
| Grade 1-2 | 5 | 342/590 | 312/532 | 0.96 (0.55-1.68) | 0.90 | 78 |
| Grade ≥ 3 | 6 | 27/635 | 23/577 | 1.14 (0.64-2.01) | 0.66 | 0 |
| Diarrhea |  |  |  |  |  |  |
| Grade 1-2 | 5 | 226/590 | 182/532 | 1.15 (0.90-1.47) | 0.26 | 47 |
| Grade ≥ 3 | 6 | 35/635 | 19/577 | 1.94 (1.08-3.47) | 0.03 | 0 |
| Fatigue |  |  |  |  |  |  |
| Grade 1-2 | 5 | 272/590 | 126/532 | 3.69 (1.99-6.85) | < 0.001 | 71 |
| Grade ≥ 3 | 6 | 25/635 | 7/577 | 2.80 (1.29-6.07) | 0.009 | 13 |
| Anorexia |  |  |  |  |  |  |
| Grade 1-2 | 5 | 310/590 | 132/532 | 4.00 (3.06-5.25) | < 0.001 | 10 |
| Grade ≥ 3 | 6 | 19/635 | 2/577 | 6.12 (1.79-21.00) | 0.004 | 0 |
| Nausea and vomiting |  |  |  |  |  |  |
| Grade 1-2 | 5 | 227/590 | 61/532 | 6.01 (2.94-12.26) | < 0.001 | 74 |
| Grade ≥ 3 | 6 | 20/635 | 4/577 | 4.10 (1.53-11.01) | 0.005 | 0 |

### Risk of bias and publication bias

As defined by the Cochrane’s manual for systematic reviews, all of included studies had a low risk of bias (S1 Fig). In addition, no publication bias for PFS and OS was found based on Begg’s test (P = 0.05 and P = 0.39, respectively; S2 Fig).

## Discussion

As the most common driver gene of lung adenocarcinoma, the status of EGFR mutation, has been gradually founded to be the most useful predictor of efficacy for EGFR-TKI over the past decade [34]. The addition of chemotherapy to EGFR-TKI as first-line treatment for EGFR-mutated NSCLC has been reevaluated to overcome or delay resistance and prolong survival time [10]. To comprehensively assess the effectiveness and toxicity of EGFR-TKI combined with chemotherapy, we systematically reviewed published randomized trials and performed a meta-analysis. The meta-analysis included eight RCTs with a combined total of 1,349 participants with EGFR-mutated NSCLC. Our results demonstrated that compared with EGFR-TKI monotherapy, the combination of first-generation EGFR-TKI and CT, especially when applying concurrent delivery of double-agents CT, significantly improve ORR (RR = 1.18, 95% CI = 1.10–1.26; p < 0.00001) and prolong PFS (HR= 0.56 (95% CI=0.50-0.64; P < 0.00001) and OS (HR=0.70 (95% CI=0.54-0.90; P=0.005), in first-line treatment for advanced NSCLC harboring activating EGFR mutation.

Growing evidence suggests that exon 19 deletions and L8585R point mutation are two different disease entities in the matter of response to TKIs and prognosis [35–38]. Kuan et al. conducted a meta-analysis of eight trials comparing EGFR-TKI with chemotherapy as first-line treatment in EGFR-mutated NSCLC [37]. Their results showed that TKI monotherapy demonstrated PFS benefit in patients with exon 19 deletions (HR=0.27, 95% CI=0.21–0.35) more than L858R (HR=0.45, 95% CI=0.35–0.58). How about the results when TKI combined with chemotherapy? In our study, the pooled HR of PFS in exon 19 deletion and L858R point mutation from five trials was 0.54 (95% CI=0.45-0.65) and 0.52 (95% CI=0.42-0.65), respectively, which were consistent. It indicated that compared with 19 deletion, the PFS of patient with L858R could be more prolonged after TKI combined with chemotherapy. This might be related to increase in ORR of L858R patient after combined chemotherapy.

Currently, first-generation EGFR-TKI monotherapy is still the mainstay of first-line treatment for EGFR-mutated NSCLC. Although the PFS can be substantially prolonged by first-generation TKI compared with platinum-based doublet chemotherapy, none of the first-generation TKIs provide an overall survival benefit revealed by several meta-analyses [39–42]. Development of new-generation TKIs or combined therapy is promising strategy to improve OS. Recent ARCHER 1050 [43] and FLAURA [44] trials have shown that second-generation dacomitinib and third-generation osimertinib, significantly prolong the OS and then both of them have been approved for first-line treatment in EGFR-mutated NSCLC [45]. As for combined strategy, adding chemotherapy to EGFR-TKI is main approach. Our meta-analysis indicated that first-generation TKI in combination with chemotherapy significantly improved the OS compared with EGFR-TKI alone (HR=0.70, 95% CI=0.54-0.90, P=0.005). Despite head-to-head RCTs are lacking in directly comparing the efficacy, first-generation EGFR-TKI combined with chemotherapy seems to prolong OS more than dacomitinib (HR=0.76, 95% CI=0.58-0.99, P=0.044) and osimertinib (HR=0.80, 95% CI=0.64-1.00, P=0.046), according to the results of HR. Based on those inspiring results, third-generation osimertinib combined with chemotherapy is speculated as a treatment strategy that could maximize the length of OS in EGFR-mutated NSCLC patients. Studies on the combination of osimertinib with chemotherapy in EGFR-mutated NSCLC, including TAKUMI and FLAURA2 trials, are currently ongoing and eagerly awaited [10].

Preclinical data indicate that the intercalated or sequential combination of EGFR-TKIs with cytotoxic agents has shown more efficacy than in the concurrent way. A possible explanation is that TKI drugs could induce the G1-phase arrest of tumor cells, which conferred a protection against the cytotoxic activity of pemetrexed [46–48]. In our subgroup analysis, however, we founded that the benefit of PFS in concurrent administration (HR=0.55, 95% CI=0.47-0.64) were consistent with in intercalated administration (HR=0.57, 95% CI=0.45-0.73), and that only concurrent administration did confer a OS benefit to patients with EGFR-mutated NSCLC (HR= 0.65, 95% CI=0.49-0.86). Our indirectly compared results could be proved by the results of NEJ005 trial, which compared concurrent versus sequential alternating gefitinib and chemotherapy in previously untreated NSCLC with sensitive EGFR mutations [49]. In addition, our subgroup analysis revealed an OS benefit in doublet chemotherapy combination group (HR: 0.67, 95% CI: 0.48-0.94), not in single-agent chemotherapy combination group (HR: 0.80, 95% CI: 0.56-1.14). But this conclusion should be applied with caution in clinical practice because only two studies adopted single-agent chemotherapy.

Adding chemotherapy to TKI also increased toxicity, notably, while increasing efficacy. Our meta-analysis indicated that most of the increased toxicities were a result of chemotherapy-induced myelosuppression and gastrointestinal toxicity, as may be expected. The incidences of serious (grade 3 or higher) hematologic toxicities from chemotherapy combination group in the meta-analysis, including leukopenia (12.1%), neutropenia (21.7%), thrombocytopenia (7.3%) and anemia (11.7%), were similar with those landmark trials in which platinum-based doublet chemotherapy were used as first-line treatment approach for the control group [50–53]. Otherwise, no significant differences were founded in terms of TKI-induced toxicities, such as any grade rash and grade 3 or higher liver dysfunction, when applying combined treatment in our meta-analysis. Meaningfully, TKI combined with chemotherapy did not significantly increase each other’s serious side effects. Therefore, the toxicities of combination therapy are manageable. Our subgroup analysis of toxicity found that compared with doublet chemotherapy, single-agent approach did not significantly increase grade 3 or higher hematologic toxicities, it may because chemotherapeutic drug in the single-agent group both adopted pemetrexed, which is generally considered to have mild toxicity [54, 55].

Admittedly, our meta-analysis has several limitations. First of all, some of results, especially in subgroup analysis, did not cover all enrolled patients due to the deficiency of detailed data, which might have an impact on the conclusion. Moreover, the outcome of OS would be confounded by the low proportion of patients in the controlled group receiving chemotherapy after experiencing progression on first-line TKI monotherapy and our study was underpowered for assessment of such effect. Thirdly, Thirdly, the subgroup analyses of different first-generation EGFR-TKI drugs, including gefitinib, erlotinib and icotinib, were not performed due to in lack of sufficient included studies. Thus, it is not clear whether the efficacies of different first-generation drugs will have differences when combined with chemotherapy. Finally, all included literatures in the meta-analysis were English language publications, which may omit other languages’ studies so as to increase the publication bias.

## Conclusions

our results demonstrate that compared with first-generation EGFR-TKI monotherapy, the combination of EGFR-TKI and chemotherapy, especially when applying concurrent delivery of platinum-based doublet chemotherapeutic drugs, significantly improve ORR and prolong PFS and OS of first-line treatment in advanced NSCLC patients harboring activating EGFR mutation, yet at the price of increasing incidence of chemotherapy-induced toxicities that is well tolerated and clinically manageable. Thus, the combination of first-generation EGFR-TKI and chemotherapy may represent a new option for first-line treatment in EGFR-mutated NSCLC. Greatly inspired by the promising results of first-generation TKI combination therapy, the results of ongoing randomized trials regarding third-generation EGFR-TKI, such as osimertinib, in combination with chemotherapy, are eagerly awaited.

## Authors contributions

QW, WXL and FX conceived the study. QW, WXL, WL and TW performed the systematic review of the literature and FX was consulted for a final decision in case of controversy. LH performed the statistical analyses. All authors contributed in interpreting the data, writing and correcting the manuscript drafts and approved the final version.

